# Analysis of genetic signatures of tumor microenvironment yields insight into mechanisms of resistance to immunotherapy

**DOI:** 10.1101/2020.03.17.980789

**Authors:** Ben Wang, Mengmeng Liu, Zhujie Ran, Xin Li, Jie Li, Yunsheng Ou

## Abstract

**Background:** Therapeutic intervention targeting immune cells have led to remarkable improvements in clinical outcomes of tumor patients. However, responses are not universal. The inflamed tumor microenvironment has been reported to correlate with response in tumor patients. However, due to the lack of appropriate experimental methods, the reason why the immunotherapeutic resistance still existed on the inflamed tumor microenvironment remains unclear.

**Materials and methods:** Here, based on integrated single-cell RNA sequencing technology, we classified tumor microenvironment into inflamed immunotherapeutic responsive and inflamed non-responsive. Then, phenotype-specific genes were identified to show mechanistic differences between distant TME phenotypes. Finally, we screened for some potential favorable TME phenotypes transformation drugs to aid current immunotherapy.

**Results:** Multiple signaling pathways were phenotypes-specific dysregulated. For example, Interleukin signaling pathways including IL-4 and IL-13 were activated in inflamed TME across multiple tumor types. PPAR signaling pathways and multiple epigenetic pathways were respectively inhibited and activated in inflamed immunotherapeutic non-responsive TME, suggesting a potential mechanism of immunotherapeutic resistance and target for therapy. We also identified some genetic markers of inflamed non-responsive or responsive TME, some of which have shown its potentials to enhance the efficacy of current immunotherapy.

**Conclusion:** These results may contribute to the mechanistic understanding of immunotherapeutic resistance and guide rational therapeutic combinations of distant targeted chemotherapy agents with immunotherapy.

## Introduction

Although immunotherapy has revolutionized tumor treatment, it still has some limitations[1]. For example, the success of adoptive cell therapy (ACT) on hematological malignancies can’t be reproduced on solid tumors[2]. The responsive rate of Immune Checkpoint inhibitors (CPI) variates by tumor type, from 45% for melanoma[3, 4] to only 12.2% for head neck squamous cancer (HNSC)[5, 6].

To better understand the reasons for these limitations, a number of studies try to investigate the phenotype of tumor microenvironment (TME) and suggesting TME phenotype (broadly categorized as being inflamed or non-inflamed) [7, 8] is a critical factor responsible for these limitations[9–13]. However, for the lack of appropriate experimental methods, a systematic understanding of how inflamed tumor microenvironment forms and why therapeutic resistance still exists on inflamed TME has been constrained. Here, to better understand the role of TME phenotypes to aid current immunotherapy, we systematically analyzed pan-cancer molecular characteristics of inflamed TME and further delved into the mechanistic differences between inflamed responsive TME and inflamed non-responsive TME. Importantly, part of our results has been supported in recent reports[14, 15].

Together, these results have a profound clinical application prospect including the identification of multiple potential immunotherapeutic targets, providing mechanistic insights into immunotherapeutic resistance in inflamed TME, and screening for some potential immunophenotypic regulation drugs to guide rational combination of chemotherapy agents with immunotherapy.

## Methods

### 1. Pan-cancer samples and clinical cohorts treated by immunotherapy

RNA sequencing data across 19 TCGA tumor types was downloaded from the Gene Expression Omnibus (GEO) database with accession number GSE62944[16]. The updated clinical data was downloaded from *TCGAbiolinks* [17–19]. A published RNA sequencing data[20] of 101 clinical tumor samples treated by anti-CTLA4 and anti-PD1 therapy was downloaded from the GEO database with accession number GSE91061. The raw count data of RNA sequencing was normalized and quantitated by the edgeR package[21].

### 2. Identifying immune cell signature from integrated single-cell RNA sequencing data

In order to analyze the TME of different tumor types and increase the diversity of non-immune cell to obtain robust immune cell markers, we applied the seurat integration pipeline[22] to integrate two single-cell RNA sequencing data sets respectively from the Puram’s HNSC cohort (GEO accession number: GSE103322)[23] and Tirosh's melanoma cohort (GSE72056)[24]. CCA algorithm[25] derived from machine learning was used to identify anchors of cells from different tumor types for purpose of unbiased single-cell data integration[26]. Annotations of immune cells referred to the original literature and cell marker database[23, 24, 27]. Immune cell genetic signatures are defined based on the following criteria: 1. The expression of signature in the identified cluster should be greater than 0.6 2. Expression in non-immune cells (myocytes, tumor cells, endothelial, Fibroblast) should be less than 0.3 3. Adjusted *P*-value<0.001 4. Log (Fold change)>0

### 3. Unsupervised clustering algorithm to determine TME subtypes of tumor samples

Immune cell markers identified in single-cell RNA sequencing analysis were used as an input for the gene set variation analysis (GSVA) algorithm[28] to calculate the immune score for each immune cell. Then, tumor samples were classified into high-immune score (inflamed), intermediate immune score and low-immune score (non-inflamed) based on the unsupervised clustering pattern. This method has been proven as an efficient way to indirectly evaluate the phenotypes of TME[29]. By using optCluster[30] to evaluate the internal and stability indexes of the seven clustering algorithms (clara, diana, hierarchical, kmeans, model, pam, sota), the optimal number and algorithm of clustering were determined. Finally, Clara algorithm and three groups were selected as the most robust clustering parameter. To avoid the unfavorable bias of confounding factors, we excluded intermediate immune score samples in further analysis.

### 4. Identification of altered signaling pathways

Differentially expressed genes (DEGs) were identified by edgeR package[21] with negative binomial distribution algorithm, *P<0.05* and log FC >1.5 were considered as statistically significant. Then, we annotated these DEGs with ClusterProfile[31] and RectomePA[32] package according to KEGG and Rectome pathway databases. Finally, gene set enrichment analysis (GSEA) was used to provide a systematic view into molecular pathway alternation[33].

### 5. Screening for potential phenotypes transformations drugs

To discover potential drugs aiding current immunotherapy, we calculated connectivity score[34] of multiple drugs to evaluate whether it is promising to promote the transformation of favorable TME phenotypes. This analysis was carried on *PharmacoGx* packages[35]。

### 6. Statistical analysis

To assess the prognostic significance of TME subtypes, we used a cox test to calculate its hazard ratio. Then, Kaplan-Meier curves and log-rank test were used to assess the differences in the 5 years’ and all years’ overall survival times between inflamed and non-inflamed subtypes. Chip-seq test and Fisher’s exact test were used to calculate *P*-value for discrete variable. *P*-value <0.05 was regarded as statistically significant.

## Results

### 1. Integration of single-cell RNA sequencing data sets

The overall design of this study was shown in Figure.1.

**Figure.1.**
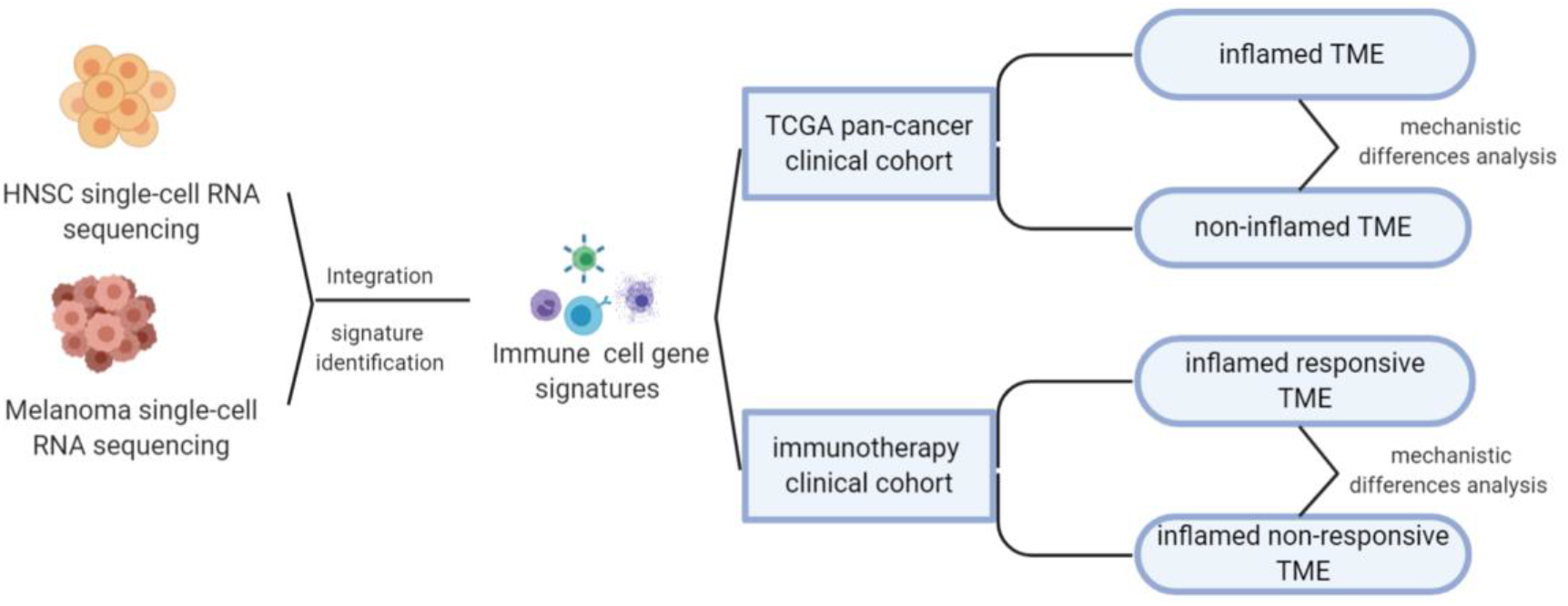
Overall design of this study.

As mentioned above, the responsive rate of immunotherapy variates by tumor type. To understand the factors that contribute to the differences in susceptibility to immunotherapy, we integrated two single-cell RNA sequencing datasets respectively from Head and Neck squamous carcinoma (HNSC) and melanoma, which were characterized by different immunotherapeutic sensitivity (approximately 45% response rate for melanoma [3, 4], significantly higher than the 12.2% of HNSC[36]).

The integration result was shown in Figure.2 A-C, tumor cells from HNSC and melanoma exhibited significant heterogeneity. Nevertheless, immune cells from different tumor types were integrated into corresponding immune cell clusters. These results suggested that immune cells from distant tumor types have a relatively similar transcriptomic pattern, which may explain the reason why immunotherapy was always accompanied by a pan-cancer therapeutic effect. The heterogeneity of immunotherapeutic efficacy across distant tumor types may be mainly derived from different tumor cells and their tumor immune microenvironment characteristics, such as immune cell composition.

**Figure.2.**
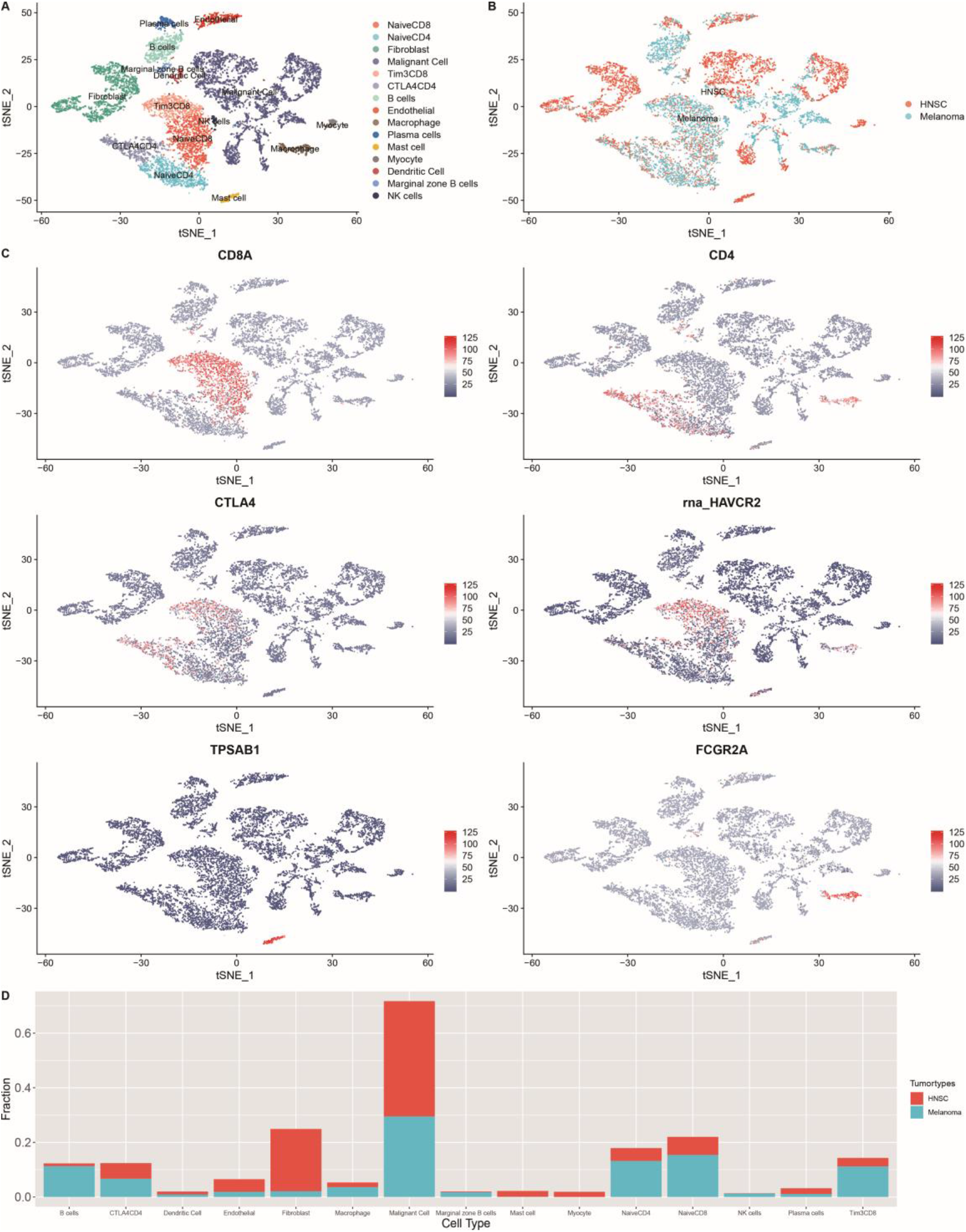
Integrated single-cell RNA sequencing analysis revealed the microenvironment heterogeneity of distant tumor types. A). The t-SNE plot displays immunological and non-immunological cells in the tumor microenvironment. Each dot represents a cell and color represents different types of cells. B). The color was coded according to tumor types. C). The expression of cell markers across different cell clusters. D). The composition of cells in HNSC and melanoma.

For instance, B cells are increasingly valued for their important role in immunotherapeutic resistance[37]. As shown in the figure.2 D, the proportion of B cells in melanoma was significantly higher than that of HNSC (P < 0.001, supplementary Table.1).

### 2. Pan-cancer prognostic significance of TME subtypes

To classify TME phenotypes across distant tumor types, immune cell gene signatures (GS) were identified in the above integrated single-cell data. Then, we classified TCGA pan-cancer samples into three TME subtypes based on the unsupervised clustering pattern of GS, each assigned as high-immune score (Inflamed), intermediate immune score, low-immune score(non-inflamed). (Figure.3 A) As shown in Figure.3 B, the proportions of TME subtypes varied greatly among different types of tumors. Next, we examined the association of this classification with overall survival time of tumor patients. Consistent with previous reports from immunohistochemistry[38], favorable prognostic roles of inflamed TME were observed in most tumor types (such as SKCM, UCEC, et al). Unexpectedly, as reported in a number of previous reports, unfavorable prognostic role of inflamed TME was also observed in some tumor types, such as LGG[39].

**Figure 3.**
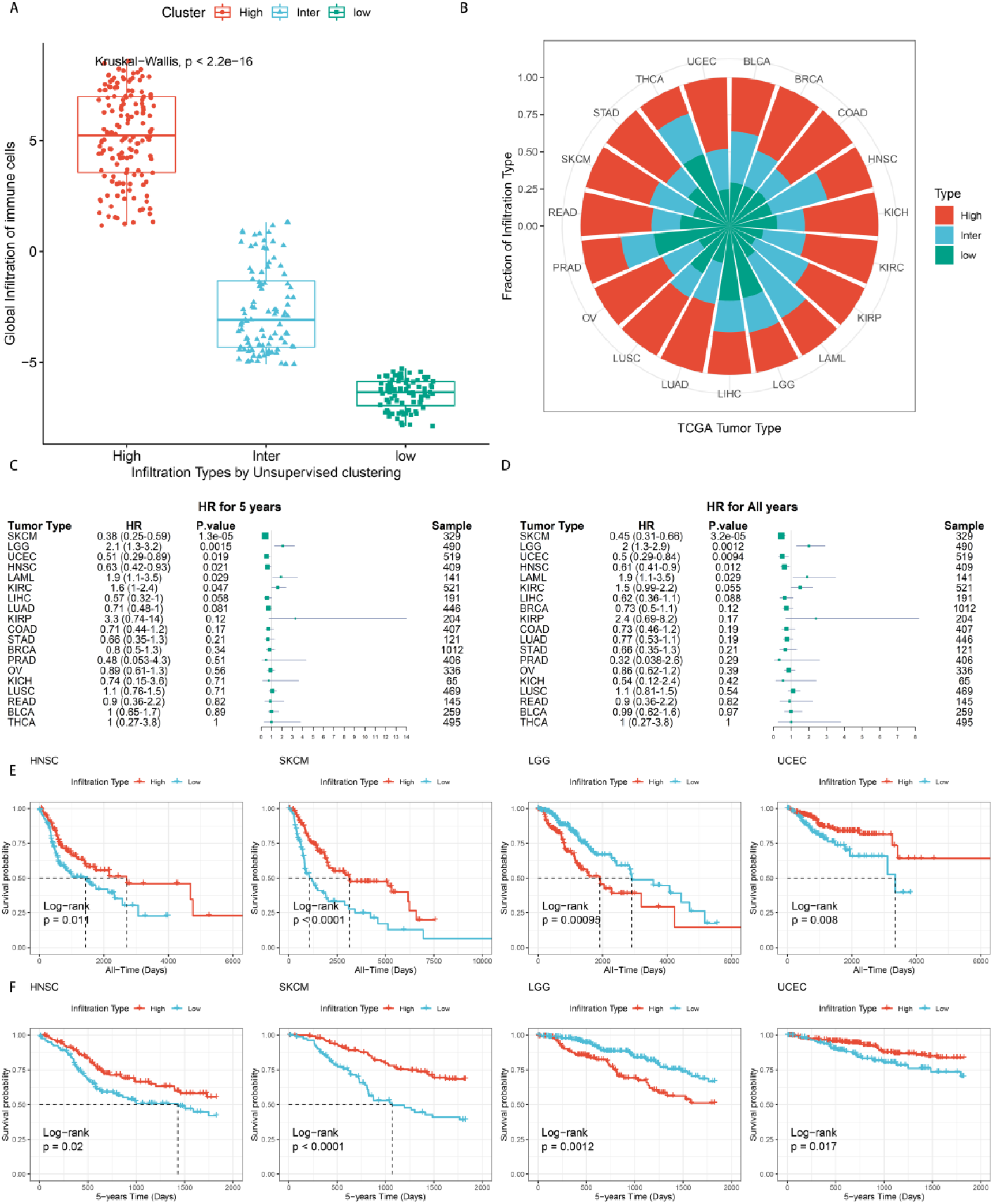
Pan-cancer prognostic role of identified TME subtypes. A). The global infiltration characteristics of distant TME subtypes. B). The proportion of TME subtypes in different cancer types. C-D). Forest plot for the association between identified TME subtypes and overall survival time of patients. E-F). All years or 5 years overall survival time of inflamed(High) and non-inflamed (Low) TME. Tumor types were represented by TCGA standard abbreviations. LAML, Acute Myeloid Leukemia; BLCA, Bladder Urothelial Carcinoma; LGG, Brain Lower Grade Glioma; BRCA, Breast invasive carcinoma; COAD, Colon adenocarcinoma; KICH, Kidney Chromophobe; KIRC, Kidney renal clear cell carcinoma; KIRP, Kidney renal papillary cell carcinoma; LIHC, Liver hepatocellular carcinoma; LUAD, Lung adenocarcinoma; LUSC, Lung squamous cell carcinoma; OV Ovarian serous cystadenocarcinoma; PRAD, Prostate adenocarcinoma; READ, Rectum adenocarcinoma; SKCM, Skin Cutaneous Melanoma; STAD, Stomach adenocarcinoma; THCA, Thyroid carcinoma; UCEC, Uterine Corpus Endometrial Carcinoma;

### 3. Molecular characteristics of inflamed or non-inflamed TME across multiple tumor types

To further investigate mechanistic differences between inflamed and non-inflamed TME, we compared gene expression profiles between inflamed and non-inflamed TME. As shown in figure.4A, up-regulated genes in non-inflamed TME related to processes such as activated GPCR signaling pathway.

**Figure.4.**
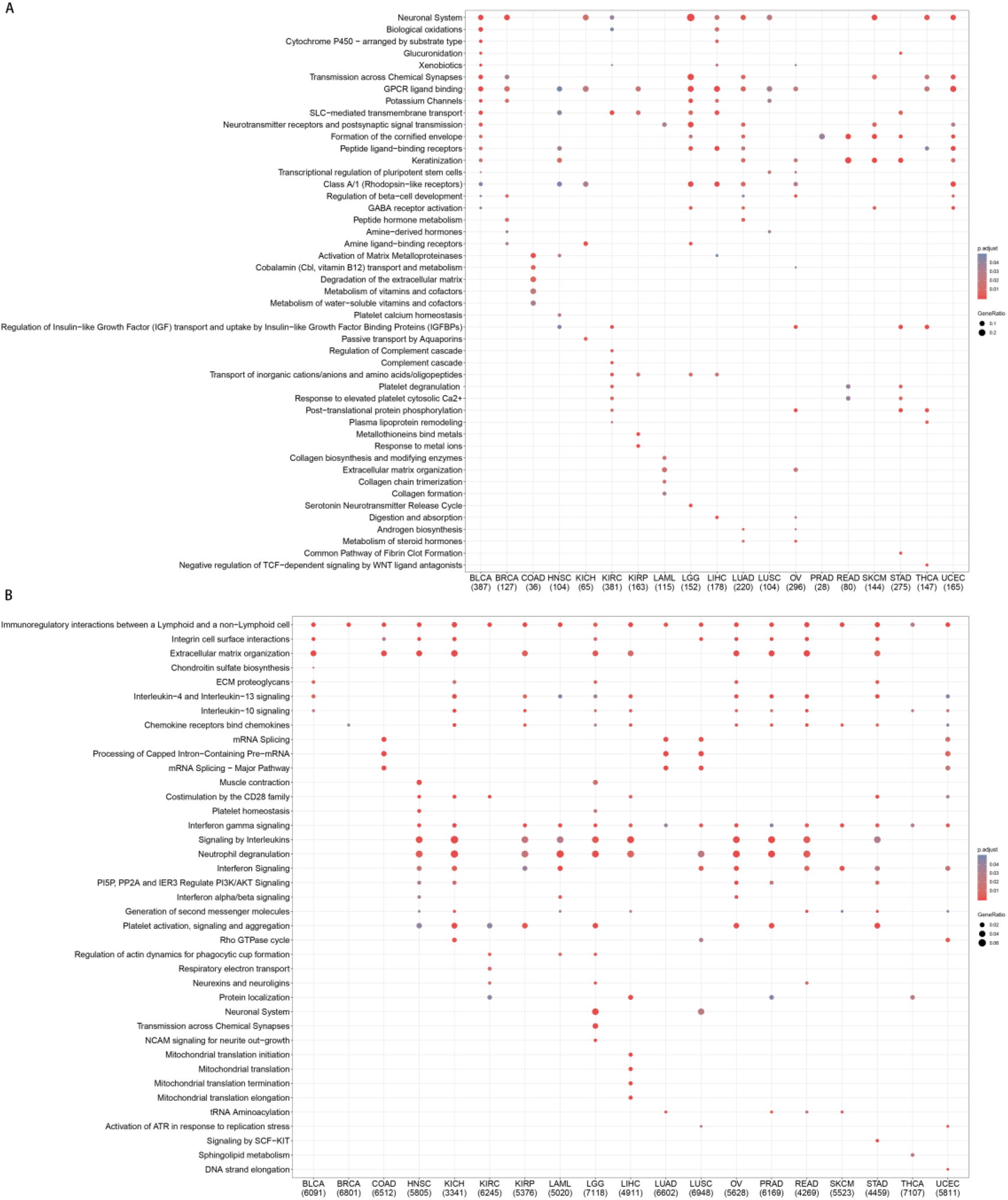
Pan-cancer gene functional annotation of TME phenotypes associated genes. A). Gene function of upregulated genes in non-inflamed TME B). The function of inflamed TME associated genes. The number under abbreviation represents the number of differently expressed genes (DEGs).

In inflamed TME (figure.4B), upregulated genes mainly related to multiple cytokine pathways, such as IFN and the interleukin-4, interleukin-13, and interleukin-10. In addition, the CD28 costimulatory molecule family associated signaling pathways, which includes PD-1, CTLA-4, was also significantly upregulated in the inflamed TME.

### 4. TME phenotypes correlated with the immunotherapeutic sensitivity

To better understand the association between TME phenotypes and the response to immunotherapy, we reproduced our classification in a published clinical cohort treated by immune checkpoint inhibitors (CPI)[40] (Figure.5 A).

**Figure.5.**
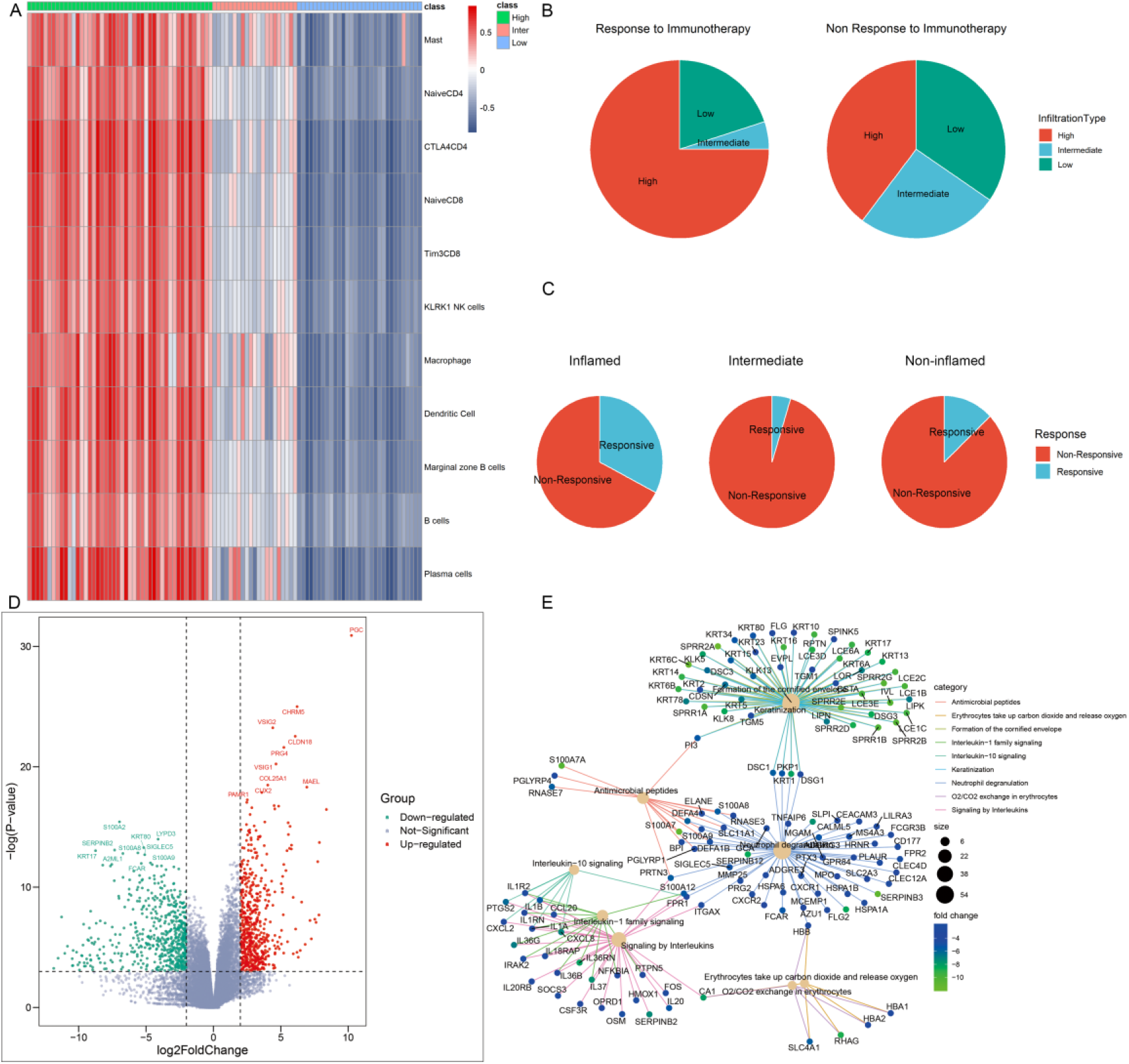
Identification of phenotype-specific genes. A). Heatmap of distant TME subtypes determined by unsupervised clustering algorithm B-C). TME subtypes correlated with the response to immunotherapy. D). The volcano plot of differently expressed genes based on an RNA sequencing analysis of inflamed responders vs inflamed non-responders. Red and green dots respectively represented upregulated-genes and down-regulated genes in inflamed responders. E). The network plot showing common genes shared by top functional terms of upregulated genes in inflamed non-responders.

As expected, inflamed tumors were most sensitive to CPI (*P*=0.015) (Figure.5 B), but only a percentage of which were responsive to CPI (Figure.5 C).

To further offer mechanistic insights into CPI resistance in inflamed TME, we identified several differentially expressed genes (DEGs) in inflamed non-responders versus inflamed responders (Figure.5 D). These genetic signatures of TME phenotype may sever as potential targets for improving current immunotherapy.

For instance, CLDN18 was the signature of inflamed responsive TME. Therapy that directly targets on CLDN18 has shown its potential to improve the efficacy of ACT in treating solid tumors[41]. On the other side, inhibiting the signature of inflamed non-responsive TME may be another promising way. Here, SIGLEC5 was significantly overexpressed in inflamed non-responders, and its family member SIGLEC15 has been proven as an efficient target to enhance anti-tumor immunity [15].

### 5. Mechanistic differences between inflamed responsive TME and inflamed non-responsive TME

Then, gene functional annotation analysis was used to understand the role of TME phenotype-specific genes. As shown in Figure.6 A, genes upregulated in inflamed and responsive tumors enriched on complement cascade and bile metabolism. GSEA analysis also confirmed multiple metabolism associated pathways were activated in this type of TME (Figure 6E). In terms of inflamed non-responsive tumors, signaling pathways, such as IL-13, IL-4, IL-10 and IL-1 cytokines related signaling pathways and oxygen exchange pathway were upregulated (Figure 5E, 6 B). These results suggested tumor hypermetabolism or activated multiple cytokines expression may confer resistance to immunotherapy.

**Figure.6.**
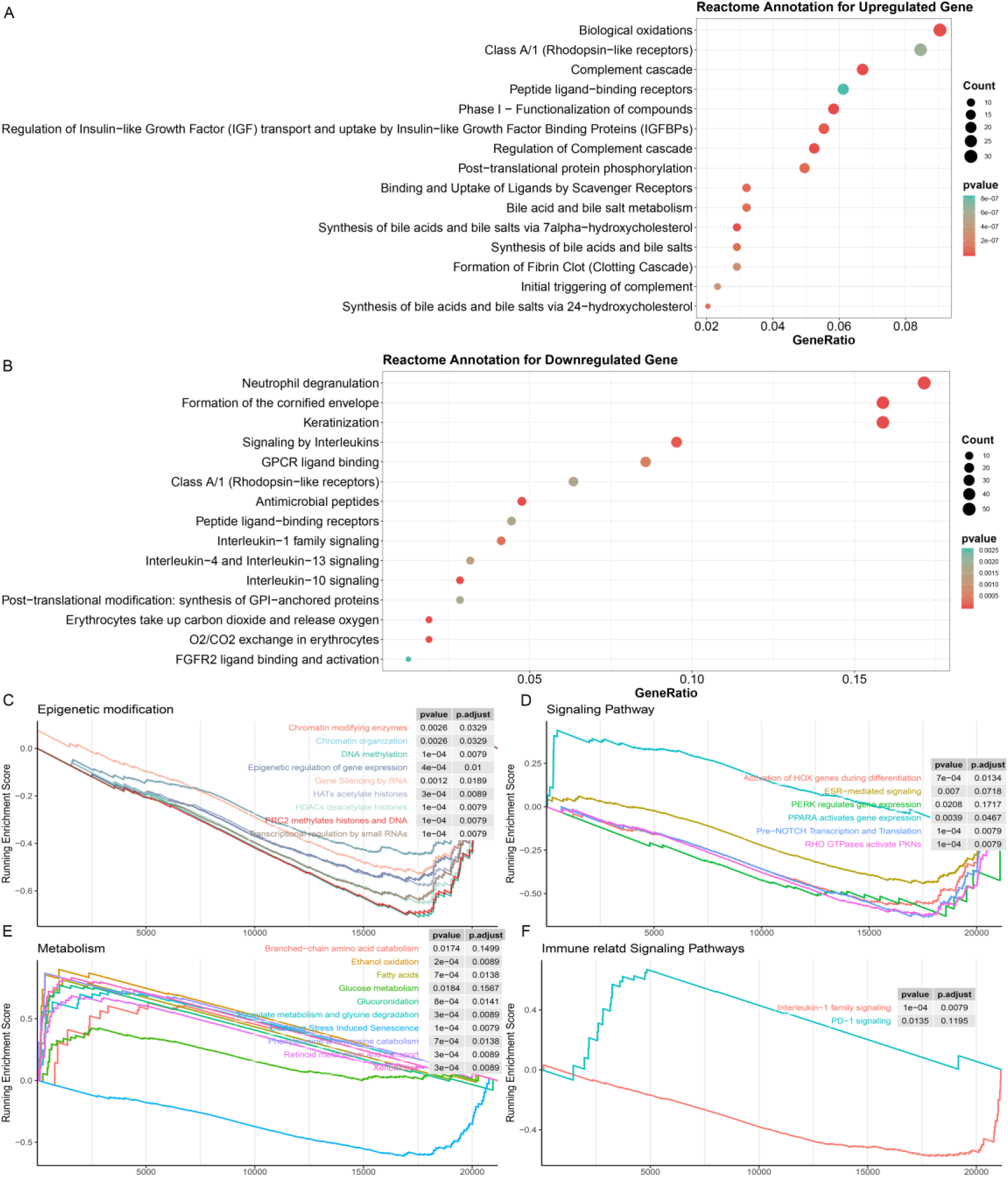
Biological processes correlated with identified DEGs in inflamed responders. A). Detailed function annotation of Up-regulated genes in inflamed responders B). Functional annotation of Down-regulated genes in inflamed responders. C-F) GSEA analysis shows four dimensions of molecular function of DEGs across inflamed responders.

Finally, for a more systematic understanding of the resistant mechanism, we applied GSEA analysis to investigate the alternation of the molecular pathways across four dimensions. (Epigenetic modification, immune or others associated signaling pathway, metabolism).

As shown in Figure.6 C, multiple epigenetic signaling pathways were activated in inflamed non-responders, suggesting a mechanism of immunotherapeutic resistance as observed by others and a potential target for reversing resistance [42, 43].

In terms of inflamed responders, multiple carcinogenesis signaling pathways, except for PPAR pathway, were inhibited (Figure 6 D, F), suggesting a mechanism of therapeutic resistance and potential target for therapy. In line with this hypothesis, recent studies illustrated that PPAR agonists appeared to improve the therapeutic sensitivity of ACT and CPI therapy[14, 44].

### 7. Screening for potential favorable TME phenotypes transformation drugs

Immunotherapy combined with chemotherapy is receiving increasing interest as a promising strategy to improve the deficiencies of current immunotherapy [45]. However, it is not completely clear how best to incorporate chemotherapy with immunotherapy. Here, we calculated the genomic connectivity score of 1288 kinds of drugs to identify potential phenotypes transformation drugs which could induce systemic favorable transcriptomic alternation, including from non-inflamed TME to inflamed TME, or from inflamed non-responsive TME to inflamed responsive TME.

Mercaptopurine (6-MP) was identified as the most promising drug, which may promote the transformation of inflamed responsive TME phenotype (Table.1). Interestingly, although some reports have shown that 6-MP can enhance the vaccine-dependent antitumor immunity[46, 47], it seems to be forgotten after that. But there are increasing interests trying to use 6-MP as a drug of immune disorders, such as autoimmune hepatitis[48], inflammatory bowel disease[49] et al. This may be since 6-mercaptopurine is widely recognized as an immunosuppressive agent, but our finding implicated that immunomodulatory may be a more accurate definition of such drugs. Our results indicated that further clinical studies were needed to assess the value of combination 6-MP with current immunotherapy.

**Table.1.** Screening for potential favorable TME phenotypes transformation drugs.

| Inflamed phenotype transformation drugs |  |  | Inflamed responsive phenotype transformation drugs |  |  |
| --- | --- | --- | --- | --- | --- |
| Drugs | Connectivity | P-Value | Drugs | Connectivity | P-Value |
| clofibrate | 0.803 | 0.021 | mercaptopurine | 0.817 | 0.028 |
| metronidazole | 0.637 | 0.002 | valdecoxib | 0.309 | 0.040 |
| zalcitabine | 0.605 | 0.028 | 5253409 | 0.292 | 0.006 |
| gabexate | 0.585 | 0.012 | astemizole | 0.290 | 0.039 |
| S-propranolol | 0.581 | 0.007 | isoconazole | 0.283 | 0.003 |
| ifenprodil | 0.580 | 0.044 | pizotifen | 0.279 | 0.011 |
| sulfapyridine | 0.578 | 0.027 | econazole | 0.278 | 0.003 |
| succinylsulfathiazole | 0.570 | 0.042 | orciprenaline | 0.267 | 0.006 |

## Discussion

Molecular stratification of TME phenotypes is paving the way for a better understanding of immunotherapeutic heterogeneity. Here, based on immune cell signatures developed from integrated single-cell RNA sequencing data, we systematically analyzed the molecular characteristics of inflamed TME across multiple cancer types and provided mechanistic insights into immunotherapeutic resistance in inflamed TME.

Part of the identified mechanistic differences has been supported by recent reports. Examples highlighted by these data including up-regulation of epigenetic-related signaling pathways and inhibited PPAR-signaling pathways are observed in inflamed non-responsive tumors, suggesting potential targets for improving the sensitivity to immunotherapy. Importantly, these results are in line with prior publications, which have provided some evidence that inhibition of epigenetic modification[50] or activation of PPAR signaling pathways[14, 44] may be promising ways to overcome therapeutic resistance to immune checkpoint blockade or ACT therapy.

These abovementioned studies also reveal the molecular characteristics of inflamed TME shared by different tumor types. These results demonstrate that inflamed TME is related to enhanced cytokines expression (interferon and IL-4, 13 and 10). Interestingly, these cytokines except for interferon are also up-regulated in inflame non-responders, suggesting a dual role of these interleukins and potential targets for therapy. These results are in line with prior published reports[51, 52]. For example, IL-10 is widely recognized as an immunosuppressive cytokine, but there is increasing evidence that it has a dual role in anti-tumor immunity. Blocking or activation of IL-10 has been proven as an efficient way to enhance anti-tumor immunity in different aspects[53, 54]. According to our results, we believe that TME phenotypes should be considered as a key factor in further study design to illuminate the remaining mysteries of IL-10 function.

In addition, our results have far-reaching clinical implications including identification of multiple potential molecular targets for developing novel immunotherapy and its combined treatment strategies. For instance, the success of ACT therapy can’t be reproduced on solid tumors due to the obstacle of its microenvironment. Therefore, rather than directly targeting on whole solid tumors, selectively targeting the inflamed and immunotherapy responsive TME may be another easier therapeutic way. As expected, this hypothesis is supported by a recent report. CLDN18, a signature of inflamed and responsive TME, has been proven as an efficient target for improving efficacy of current ACT therapy on solid tumors[41].

Except for targeting on inflamed and responsive TME, examples highlighted by our data also include inhibiting the signature of inflamed non-responsive TME to reverse therapeutic resistance. For example, SIGLEC15, a signature of inflamed and non-responsive TME, has shown its power in blocking immune escape. Interestingly, the enhancement effect of anti-tumor immunity is independent of the PD-1/PD-L1 axis, suggesting that it may be an ideal target to aid current anti-PD-1 therapy[15].

Finally, based on a genomic-drugs perturbation database, we identify some drugs which are promising for promoting the transformation from unfavorable TME phenotype to favorable one.

In conclusion, our result provided an important view for understanding how inflamed TME and inflamed resistant TME forms. This evidence has important clinical implications and may help guide rational therapeutic combinations of distant chemotherapy agents with immunotherapy depending on the desired treatment effect.

## Acknowledgments

Thanks to everyone who pushed the boundaries of human knowledge

